# BTK Autoinhibition Analyzed by High-Throughput SH2 Domain Swaps

**DOI:** 10.1101/2025.02.06.636897

**Authors:** Timothy J. Eisen, Sam Ghaffari-Kashani, Chien-Lun Hung, Jay T. Groves, John Kuriyan

## Abstract

BTK, a Tec-family tyrosine kinase, resembles the Src and Abl kinases in that an SH2-SH3 module regulates the activity of the kinase domain, principally through an inhibitory interaction between the SH3 and kinase domains. In Src kinases, phosphorylation of a C-terminal tail latches the SH2 domain onto the kinase domain, positioning the SH3 domain in an inhibitory conformation; in Abl, interaction between the kinase domain and a myristoyl group on the N-terminal segment provides the same latching function. The structure of autoinhibited BTK resembles that of the Src and Abl kinases, but BTK lacks an SH2-kinase latch. To assess autoinhibition in BTK, we generated hundreds of chimeric BTK molecules and measured their fitness using a high-throughput assay in T and B cells. Surprisingly, many SH2 domains increased fitness when substituted into BTK. Analysis of one set of chimeric proteins demonstrated that the increase in fitness stems from the ability of the substituted SH2 domains to disrupt BTK autoinhibition while maintaining phosphotyrosine targeting. These results reveal the importance of distributed interactions between the SH2 and kinase domains of BTK in stabilizing the inhibitory conformation, and suggest that the specialized latching mechanisms in Src and Abl kinases may be later evolutionary refinements.

## Manuscript Text

The discovery of the SH2 domain by Pawson and colleagues^1^ revolutionized our understanding of cellular signaling. Inherent to the understanding of SH2 function was the idea of modularity: that the function of phosphotyrosine binding and intracellular targeting, conferred by the SH2 domain, could be transferred to any protein that needed it. The proliferation of SH2 domains supports the notion that these domains are modular: there are over 59,000 catalogued SH2 domains across eukarya^2^.

Phosphotyrosine recognition is only one function of the SH2 domain. In tyrosine kinases such as c-Src, c-Abl, and the Tec kinases, including Bruton’s Tyrosine Kinase (BTK), the SH2 domain is found within a “Src module”—an ancient structural unit that consists of an SH3 domain, an SH2 domain and a tyrosine kinase domain. In the Src family of kinases, the autoinhibited conformation of the Src module is stabilized by a latch, provided by a phosphorylated tyrosine in the C-terminal tail, that interacts with the SH2 domain^3,4^. The latch anchors the SH2 domain next to the C-terminal lobe of the kinase and positions the SH2-kinase linker so that it can serve as a docking site for the SH3 domain. Interaction of the SH3 domain with the N-terminal lobe of the kinase stabilizes the inactive conformation of the catalytic center^5^.The absence of the SH2 latch in the viral oncoprotein v-Src accounts for its constitutive activity^6^. In c-Abl, an N-terminal myristoyl group provides the latching function through an allosteric mechanism^7^. In the oncogenic BCR-ABL fusion protein, the myristoylation site is absent, contributing to uncontrolled kinase activity.

The Src module of the Tec-kinase BTK adopts an assembled conformation that is very similar to that of c-Src and c-Abl, as shown by crystallography and cryo-electron microscopy^8,9^, but has no obvious latching mechanism. As in Src and Abl kinases, a key autoinhibitory interface in BTK is between the SH3 domain and the kinase domain. The interface between the SH2 domain and the kinase domain is not tightly packed, suggesting that this interface does not contribute to the stability of the autoinhibited conformation in BTK. Despite this, the SH2 domain is involved in autoinhibition. A mutation in the C-terminal segment of the kinase domain, near the SH2 domain (D656K), activates the kinase^10^ and a mutation in the SH2 domain (T316A) confers drug resistance^11^.

We developed a high-throughput assay for BTK function that allows us to test the fitness of hundreds of chimeric BTK molecules with different SH2 domains (328 SH2 domains in total, Figures 1A–B and S1A). These SH2 domains were derived from vertebrate Tec kinases, human SH2-containing proteins, SH2 domains generated using ancestral sequence reconstruction, and control SH2 domains with mutations that disrupt phosphotyrosine binding. The fitness of each SH2–BTK chimera was tested using a cellular assay for BTK function that we developed previously^12^ (Figure S1B). This assay uses the ability of BTK or variants to induce expression of CD69, a surface-expressed glycoprotein, in Jurkat T cells lacking ITK or Ramos B cells lacking BTK. This ability is quantified by a fitness score, defined as the base-10 logarithm of the ratio of RNA-seq counts for a particular variant in the CD69-enriched fraction and the corresponding counts in the input fraction. Fitness scores are normalized such that human BTK has a fitness score of 0. Fitness scores agreed well among biological replicates (Figure S1C, Pearson R ≥ 0.65).

**Figure 1.**
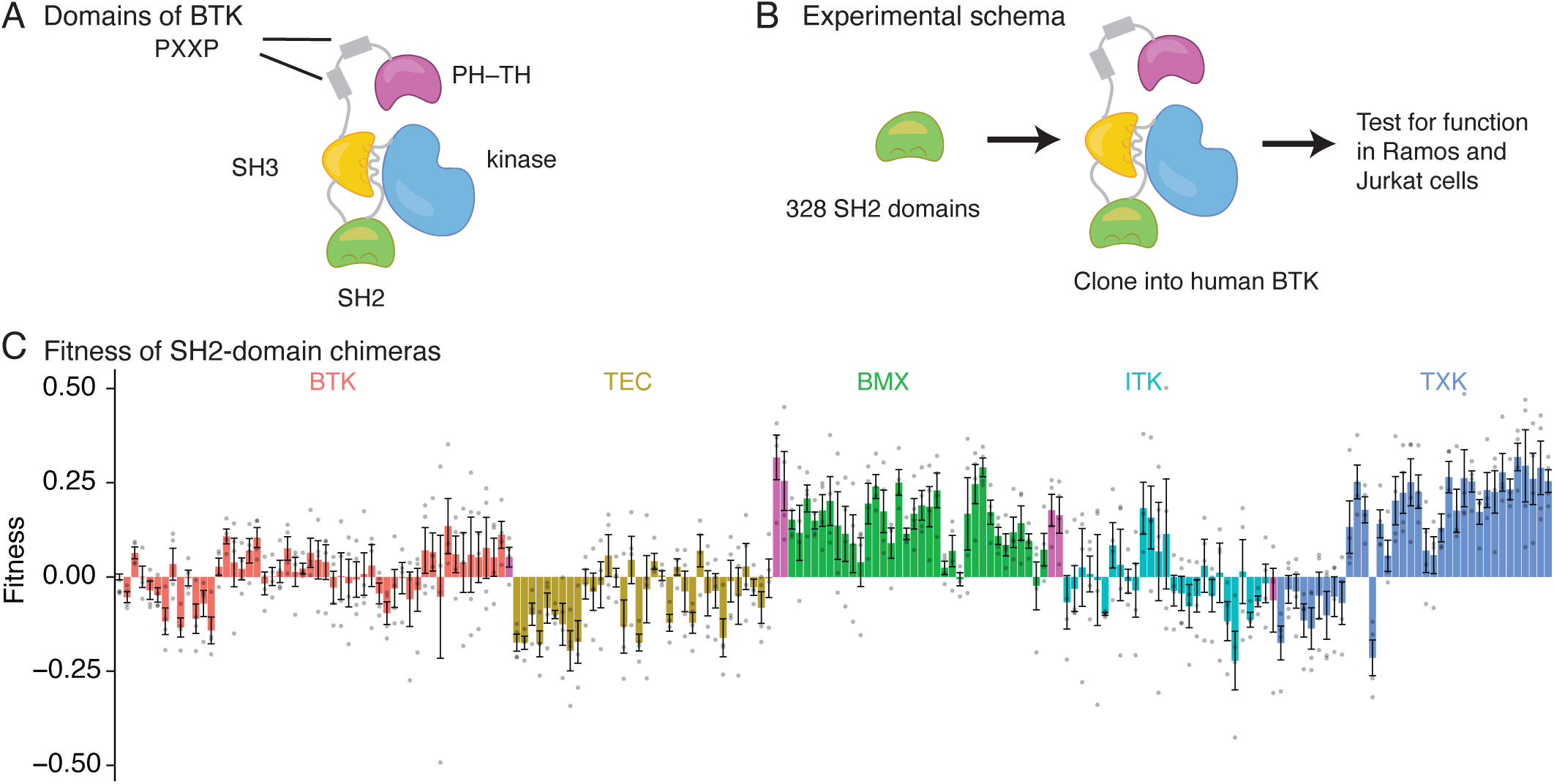
SH2-domain substitutions increase fitness in BTK. a. A schematic diagram of BTK, with all domains labeled. b. Outline of the experiment. c. Fitness scores for the 188 SH2-domain chimeras corresponding to Tec kinase or ancestral SH2 domains. SH2 domains are ordered by their sequence relationships. The fitness bars, measured using the Jurkat assay, are colored by the human Tec kinase to which they are most closely related, requiring at least 75% sequence identity to display that color (sequences without 75% sequence identity to a human Tec kinase are colored purple). Error bars represent standard error of the mean and points are the individual replicate values.

We implemented both the Jurkat and Ramos adaptations of the assay in this study, focusing first on the Jurkat data because BTK kinase activity is required for BTK to substitute for ITK in ITK^−/−^ Jurkat cells^12^. In contrast, BTK kinase activity is dispensable for signaling in Ramos cells^12,13^. Instead, Ramos cells require kinase-independent scaffolding functions of BTK, which are also important for its role in some cancers^14,15^.

Disruption of SH2 targeting by mutation of a key phosphotyrosine-binding residue (Arg 307 in BTK^16^) reduces the fitness of human BTK and almost all chimeras for which this mutation was tested (57 out of 62, Figure S1D). Thus, as expected, phosphotyrosine binding is an important function of the SH2 domain in Tec kinases. Strikingly, many SH2-domain substitutions resulted in an *increase* in fitness with respect to human BTK (Figure 1C and Table S1). Of 249 SH2-chimeras in the library (excluding controls), half (51%, or 128 domains) had fitness scores greater than 0. BTK chimeras with Tec-kinase SH2 domains (188 domains) cluster into groups that increase or decrease fitness similarly, depending on the Tec kinase that the SH2 domain is derived from (Figure 1C). Despite some measurement noise, it is clear that SH2 domains from two classes, corresponding to the BMX and TXK domains, increase fitness when substituted into human BTK. A plausible explanation for these fitness increases is that the autoinhibitory function of the SH2 domain is less readily transferrable than its phosphotyrosine-binding function, leading to activation of BTK by substitution with non-cognate SH2 domains. Similar results were observed in Ramos cells, which do not require the kinase activity of BTK (Figure S2A–C).

To determine how different SH2-domains increase fitness, we studied the BMX group (Figure 2A). We used ancestral-sequence reconstruction to infer the possible sequence of a common BMX ancestor in order to focus our attention on small sets of mutations that modulate fitness (Figure S3A–B)^17,18^. Eight reconstructed ancestral SH2 domains within the BMX group (labeled BMX-A through BMX-H) progressively increase fitness when substituted into human BTK (Figures 2B). The BTK chimera with the BMX-A SH2 domain (with 50% identity to the human BTK SH2 domain), is neutral, and behaves similarly to human BTK. The BTK chimera with the BMX-H SH2 domain is gain-of-function. Between BMX-A and BMX-H, there are four substitutions in the SH2 domain: N285D, N289S, R310S, and A312V. Strikingly, these four substitutions are located at the interface between the SH2 and kinase domains seen in the Btk crystal structure (PDB code: 4XI2), indicating that disruption of interactions between the SH2 and kinase domains increases fitness (Figure 2C). One residue, His 285 in human BTK, is an asparagine in BMX-A, a neutral BTK chimera, and an aspartate in BMX-H, a gain-of-function chimera. Asp 285 in BMX-H is located next to a set of negatively charged residues on the adjacent kinase domain and next to Tyr 223, a conserved residue in the SH3 domain that activates BTK when phosphorylated^19^. The location of Tyr 223 near other negatively charged residues, including Asp 285 in the gain-of-function chimera, suggests that pTyr 223 activates BTK by increasing electrostatic repulsion (Figure 2D). As expected, the c-Abl and c-Src SH2 domains, which require latching for autoinhibition, increase fitness when substituted into BTK (Figure S3C). Substitution by more distantly related SH2 domains tends to decrease fitness in general.

**Figure 2.**
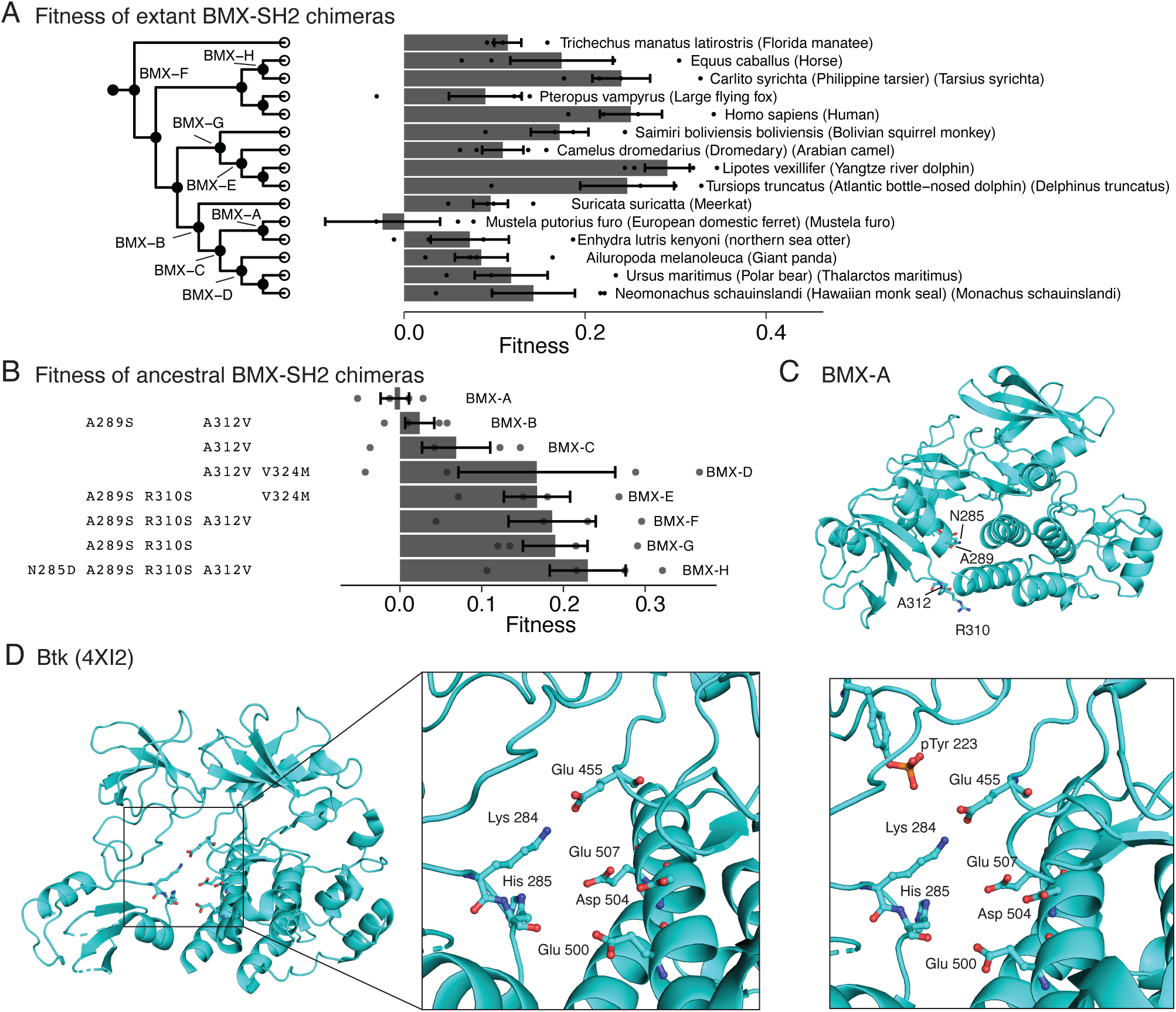
Ancestral-sequence reconstruction charts fitness progression in the BMX Lineage. a. Fitness scores from the BMX group of SH2 sequences. Each sequence is shown with its Latin name and common name. Positions correspond to sequence relationships based on a phylogenetic gene tree (left), in which ancestors that were reconstructed are labeled. Error bars represent standard error of the mean. One point was removed from the *Mustela furo* bar at x = –0.2. b. The eight ancestral-reconstructed sequences from (a), arranged by order of increasing fitness. The substitutions to the BMX-A sequence are shown at the left. c. An Alphafold^20^ model of the Src module of the BMX-A sequence. The four residues (Asn 285, Ala 289, Ala 312, Arg 310) substituted in the BMX-H sequence are shown as a stick representation. d. (left and middle) Negatively charged residues that interact with His 285, shown on the crystal structure of the mouse Btk Src module (PDB: 4XI2^Ref. 8^). (right) The inset is shown with Tyr 223 (the site of autophosphorylation) modeled as a phosphotyrosine.

To explore further the role of the SH2-kinase domain interface in autoinhibition, we examined the same interface, but from the perspective of mutations in the kinase domain, rather than the SH2 domain. We mutated helix αI^kinase^, the C-terminal helix in the kinase domain (Figure S4A), which interacts with the SH2 domain in the crystal structure of Btk^8^. We created a library of αI^kinase^ sequences from 118 BTK orthologs and Tec kinases from the jawed vertebrates, and swapped these sequences for αI^kinase^ in human BTK (Figure S4B; fitness scores were reproducible, with R ≥ 0.87 for n = 236 proteins and n = 4 biological replicates, Figure S4C). In keeping with a previous study^10^, substitution of Asp 656 increased fitness (Figure S4D). Fitness scores for this library and the SH2 library agree well between Jurkat and Ramos cells (Figure S4E)^12,13^.

We sought αI^kinase^ sequences that modulate fitness by stabilizing (rather than disrupting) the inactive conformation, reasoning that such sequences would provide insight into the structural features that support autoinhibition (Figure 3A). The challenge of searching for αI^kinase^ sequences that stabilize the autoinhibited conformation is that in addition to such sequences, sequences that destabilize the protein are also expected to decrease fitness. To distinguish between these mechanisms, we measured the fitness of αI^kinase^ variants in two genetic backgrounds: BTK and the BMX-H gain-of-function SH2 chimera (Figure 3B). We developed a barcoding approach to simultaneously determine the identity of the SH2 domain and the αI^kinase^ variant (Figure S5A).

**Figure 3.**
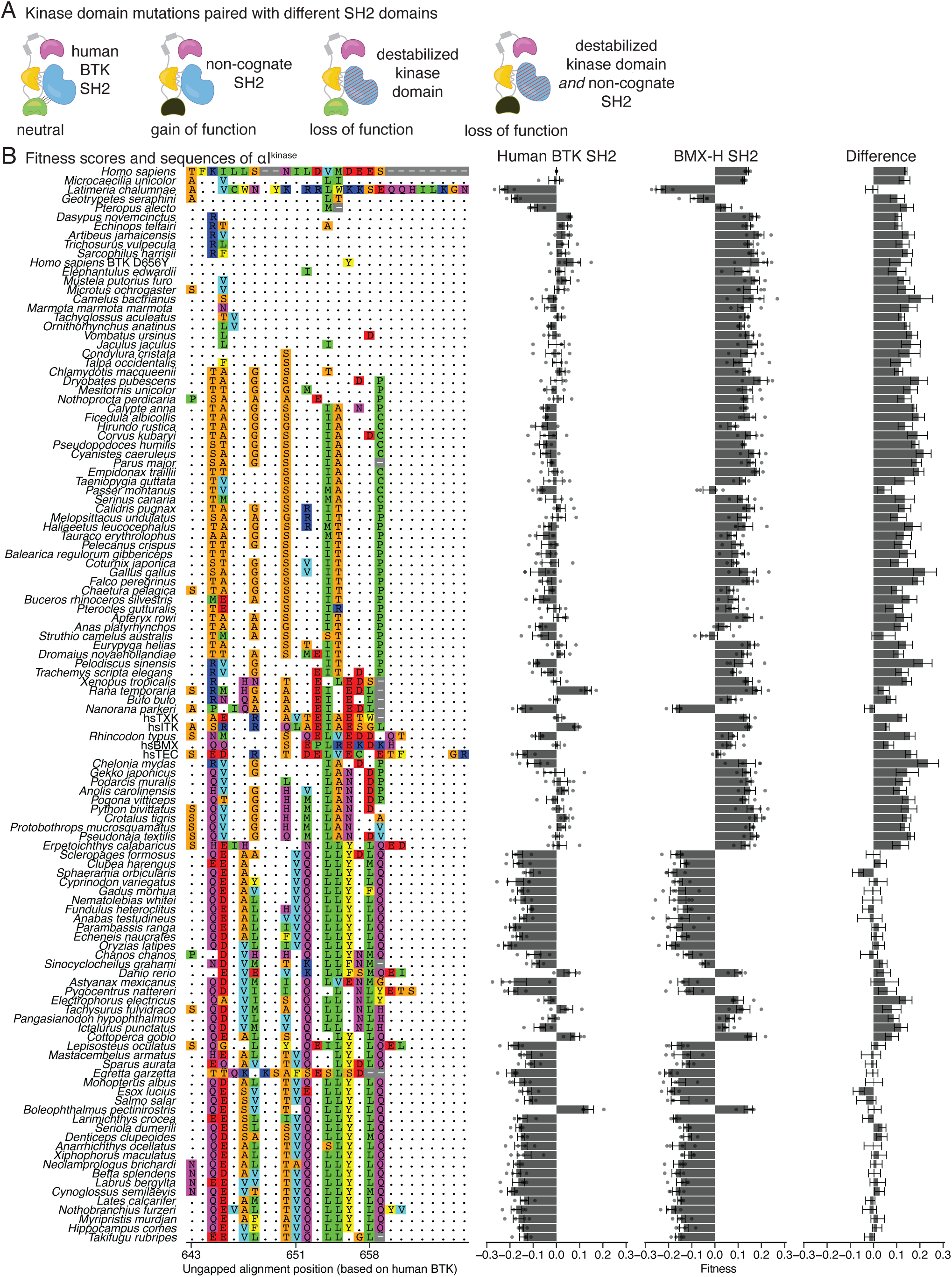
Screening αI^kinase^ substitutions. a. Schematic representation of the effect of combining a destabilizing mutation in the kinase domain with a non-cognate SH2 domain. b. The 118 αI^kinase^ sequences are shown along with a multiple sequence alignment and fitness scores. The fitness scores are shown for the human BTK SH2 genetic background, the BMX-H SH2 genetic background, and the difference between these two backgrounds (BTK values subtracted from the BMX-H values). Error bars represent standard error of the mean. For the difference values, error bars are determined using the standard error propagation formula.

An αI^kinase^ variant that destabilizes the kinase domain should reduce fitness in both BTK and the BMX-H SH2 chimera. In contrast, an αI^kinase^ variant that reduces fitness by making interactions that stabilize the BTK SH2-kinase interface should reduce fitness in BTK, but have little or no effect in the BMX-H SH2 chimera, which has a different SH2 domain. Thus, αI^kinase^ variants that display reduced fitness in both the BTK and BMX-H SH2 genetic backgrounds are likely to be less stable. αI^kinase^ variants that reduce the fitness of BTK, but not the BMX-H chimera, are likely to exhibit a greater degree of autoinhibition.

As expected, many αI^kinase^ sequences exhibit similar negative fitness scores in the human BTK and BMX-H backgrounds, consistent with destabilization (Figure 3B and 4A). Ile 651 in αI^kinase^ forms a hydrophobic packing interaction with the adjacent helix, αG^kinase^, in the C-lobe helical bundle (Figure S5B). Some of the instability arising from αI^kinase^ variants can be traced to substitution of Ile 651 by the smaller valine residue (compare e.g. *Salmo salar*, arctic salmon, and *Boleophthalmus pectinirostris*, great blue spotted mudskipper, Figure 3B).

**Figure 4.**
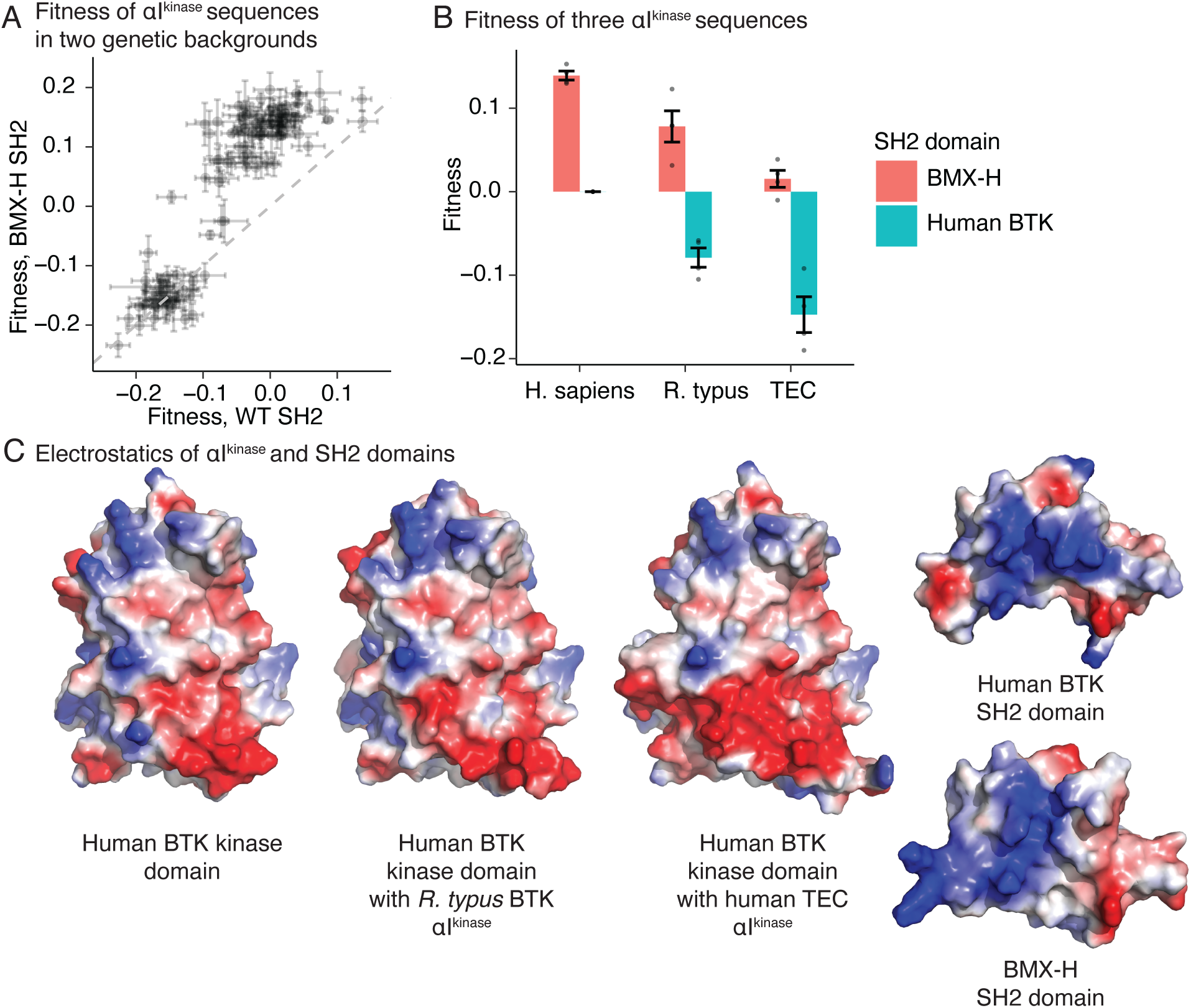
Two αI^kinase^ substitutions decrease fitness consistent with enhanced autoinhibition. a. Fitness scores from 118 αI^kinase^ sequences shown in two different genetic backgrounds. Error bars represent standard error of the mean. b. Fitness scores from three αI^kinase^ sequences (x axis) shown in two genetic backgrounds (key). c. Alphafold^20^ model of the human BTK protein, the αI^kinase^ sequence of *R. typus* in the human BTK kinase domain, and the αI^kinase^ sequence of TEC in the human BTK kinase domain. The kinase-domain interacting interfaces of the human BTK and BMX-H SH2 domains are shown at the right. Coloration is based on charge, with positively charged regions in blue and negatively charged regions in red.

We found two αI^kinase^ sequences, derived from human TEC and from *Rhincodon typus* (whale shark) BTK, that decreased fitness in human BTK but not in the BMX-H SH2 chimera (Figure 4B). Structures of the human BTK kinase domain with *R. typus* or TEC αI^kinase^, generated by Alphafold^20^, show an increase in net negative charge on αI^kinase^ that parallels their relative fitness (Figure 4C and Figure S5C). The kinase-binding region of the BTK SH2 domain has higher positive electrostatic potential with respect to the corresponding region of the BMX-H SH2 domain (Figure 4C). Thus, the two αI^kinase^ variants are consistent with a mechanism in which enhanced electrostatic interaction between the SH2 and kinase-domain interfaces decreases fitness.

In conclusion, we examined the effect of replacing the SH2 domain of BTK with hundreds of SH2 domains from other proteins. As expected from the idea that SH2 domains are modular units, most of the chimeric BTK proteins that we tested are functional substitutes for human BTK. Surprisingly, we found that many of the chimeras exhibited *increased* fitness with respect to human BTK. For one set of chimeric proteins, we traced these fitness increases to disruption of the SH2-kinase domain interface seen in the structure of inactive Btk^8^, demonstrating that the SH2 domain is autoinhibitory even though BTK lacks an obvious latching mechanism. The nature of the autoinhibitory interface is predominately electrostatic, with the negatively charged residues in the C-terminal helix of the kinase domain (αI^kinase^) located near positively charged residues in the phosphotyrosine-binding site of the SH2 domain. Indeed, phosphopeptide binding and SH2-kinase docking appear mutually exclusive^10^. These results suggest that in a prototypical Src module without a latching mechanism, distributed interactions between the SH2 domain and the kinase domain may have sufficed to suppress kinase activity and impede the targeting function of the SH2 domain until released by activation of the kinase domain. Kinases with this simple switching mechanism are seen in the choanoflagellates, and therefore arose before the establishment of the metazoan lineage^21^ and such kinases have survived to generate some of the most important signaling proteins in humans.

## Acknowledgements

We thank J. Paul for many helpful discussions and inspiration for the epistasis experiments. We also thank S. Choi, S. Muratcioğlu, and other members of the Kuriyan and Groves laboratories for helpful discussions. This work was greatly assisted by the Cell and Tissue Analysis Facility, the Vincent J. Coates Genomics Sequencing Laboratory, and the Cancer Research Laboratory Flow Cytometry Facility. T.J.E. is a Damon Runyon Fellow supported by the Damon Runyon Cancer Research Foundation (DRG-2429–21). This work was supported by the National Institutes of Health (P01 A1091580).

## Data and materials availability

Raw fastq files and processed files are available on DRYAD. Code for generating saturation-mutagenesis libraries and analyzing them is available on GitHub (https://github.com/timeisen/MutagenesisPlotCode and https://github.com/timeisen/SH2s). DNA libraries, plasmids, and cell lines produced in this work are available upon request.

## Methods

### Ancestral Sequence Reconstruction

All 59,304 SH2 domain sequences in the Protein Families (PFAM) database were downloaded as a multiple sequence alignment. These domains align to the human BTK SH2 domain, corresponding to residues 281–362. After de-duplication, this set contained 23,121 SH2 domains, of which 156 were derived from human proteins. This set of human SH2 domains was filtered to include only the 66 SH2 domains that had at least 25% pairwise identity to the human BTK SH2 domain. This set was then divided into Tec-kinase SH2 domains (those with ≥ 40% identity to the human BTK SH2 domain) and more distantly related SH2 domains (those with identity to the human BTK SH2 domain between 25% and 40%). The six human Tec SH2 domains consisted of: TEC, ITK, TXK, BMX, Q5JY90 (an isoform of BTK), and BTK.

The six human Tec SH2 domains were then used to mine the original set of unique PFAM domains for an additional set of 84 non-human Tec-kinase domains. These domains were selected from other organisms by progressively choosing 14 domains for each Tec-kinase sequence with increasing pairwise identity from the human Tec-kinase domain, excluding any domains with a single mismatch to the human domain. The resulting set of 90 domains included 15 domains for each of the 6 Tec-kinase SH2 domains.

These Tec-kinase domains from humans and other organisms were used for ancestral sequence reconstruction. (One domain, A0A452S617, the Q5JY90 BTK isoform from the American black bear, was removed from the ancestral sequences reconstruction because of low quality). First, a maximum-likelihood tree was generated using Practical Alignment using Saté and TrAnsitivity algorithm^22^. Then the PAML software suite (version 4.8a, August 2014,^23^) was used to generate ancestral sequences. To perform this reconstruction, we implemented both marginal^24^ and joint^25^ reconstruction of the ancestral sequence. Marginal reconstruction is optimizing the state at the current node that maximizes the probably over the entire tree. Joint reconstruction is optimizing all states simultaneously to maximize the probability over all nodes. The model used the SWISSPROT probabilities of exchanging amino acids^26^.

PAML reconstruction relies on all sequences in the alignment and cannot determine ancestors from sequence alignments with gaps. We reasoned that regions with gaps would generally be variable regions that could be quite important for targeting specificity or kinase-domain interaction and chose two methods to reconstruct these gap sequences. In the first method, any position that contained at least one gap in the alignment was removed from the alignment entirely, resulting in all sequences aligning to the shortest sequence. In the second method, residues were included at all gaps, interpolating from all other sequences in the alignment. This method aligns all domains to the longest sequence. We implemented both methods to reconstruct ancestral nodes. This analysis resulted in 114 ancestral nodes. In Table S1, a prime (′) denotes sequences that were constructed from the longest domain and those without a prime were constructed from the shortest domain. The resulting nodes were combined with the extant Tec (90 domains) and more distant domains (60 sequences) for a total of 264 domains. This set of domains was further filtered to remove ancestral nodes from the Q5JY90 lineage (9 nodes) and domains longer than 88 amino acids (6 sequences) for a final set of 249 domains. Extant domains corresponding to Q5JY90 were excluded from Figure 1C because they have a large truncation in the SH2 domain and decrease fitness.

Control sequences were added to this set to determine the effect of disrupting the phosphotyrosine binding residue (R307K substitutions) or of completely removing BTK by adding a stop codon (R307X). We reasoned that these controls would only be informative for sequences similar to the wild type-human BTK sequence, as very distant domains, even if they increased fitness, would be more difficult to interpret the effects of phosphotyrosine targeting. Any domain of the set of 249 (including ancestral domains) with a pairwise identity greater than 60% to the human BTK SH2 domain (62 domains) was included with the R307K mutation. A subset of these 62 domains with pairwise identity to the human BTK SH2 domain of greater than 95% (17 domains) were also mutated to contain a stop codon at position 307 to determine the baseline fitness in the assay. All 79 control sequences were included with the original 249 sequences for a total of 328 SH2 domain sequences. Figures 1C and S3C, which paired the fitness data with a phylogenetic tree, were constructed using the ggtree plotting software^27^.

### Reverse Translation of Sequences

Nucleotide sequences were constructed from each of the 328 protein sequences by generating four synonymous nucleotide sequences for each protein sequence. If an amino acid in a protein sequence was identical to that of BTK, the human BTK codon was used at that position.

Otherwise, a codon was chosen at random, excluding sequences that contained the BsaI recognition sequence. 31 additional codon variants of the wild-type human BTK SH2 sequence were included by choosing 5 codon positions at random and substituting a synonymous codon. In the αI^kinase^ library, 44 synonymous wild-type sequences were included. Handles allowing for BsaI digestion were then included at each end (left handle: 5′ CAGGCATGGTCTCATGAG 3′; right handle 5′ CAGCAGAGACCCGATTGG 3′). Because the resulting nucleotide sequences were different lengths depending on the length of the SH2 domain, sequences were padded with additional sequence from the PhiX genome for a final length of 300 nt. Sequences were ordered from Twist Biosciences as oligo pools.

### High-throughput Cloning

The pooled oligos of SH2 domain sequences were amplified by 15 cycles of PCR using primers that annealed to the left and right constant sequences (oligos #272 and #273, Table S2). BTK constant sequences corresponding to gene segments 5′ of the SH2 region and 3′ of the SH2 region were also amplified in separate reactions using a BTK template in which endogenous BsaI restriction sites were removed by replacing them with synonymous codons. These constant segments were amplified with oligos #274 and #275 for the 5′ segment, and oligos #276U and #277 for the 3′ segment (Table S2). The left oligo of the 5′ segment encodes a XmaI restriction site while the right oligo of the 3′ segment encodes a BamHI restriction site.

The three pieces of BTK (corresponding to the amplified pool of SH2 domains, the 5′ constant BTK segment, and the 3′ constant BTK segment) were agarose-gel purified and combined in a 50 µL reaction containing 2 µL of BsaI HF (New England Biolabs), 2 µL of T4 DNA ligase (New England Biolabs), 5 µL of 10x T4 DNA ligase buffer (New England Biolabs) and 0.5 µL of DpnI (New England Biolabs). The DpnI was intended to remove any contaminating uncut vector, which was the template in the PCR of the constant sequences. This ligase mixture was incubated for two hours at 37 °C and then overnight cycling between 12 °C for 10s and 37 °C for 10 min. The reaction was run on a 1% agarose gel and the fully ligated product (∼2 kb) was excised and purified.

The assembled BTK sequence was digested with BamHI HF (New England Biolabs) and XmaI (New England Biolabs) in a 30 µL reaction containing 3 µL of CutSmart buffer (New England Biolabs). The digested sequence was ligated into a vector backbone (Addgene #21373) containing an internal ribosome entry site that had been digested with BamHI, XmaI, and EcoRI (EcoRI digests the BTK gene, and is included when preparing the vector backbone to decrease contamination from uncut human BTK). The ligation reaction used a 5 to 1 insert to vector ratio and 100 ng of vector. After 1 h at room temperature, the ligation reaction was purified using the Zymo Clean and Concentrator kit (Zymo Research) according to the manufacturer’s instructions and DNA was eluted in 6 µL. 3 µL of the elution was electroporated into Endura electrocompetent cells (Lucigen) according to the manufacturer’s instructions. Cells were recovered in 1 mL of Recovery Media (Lucigen) for 1 h at 30 °C and diluted to 50 mL in Luria Broth. 50 µL of this dilution were plated as a diagnostic to gauge transformation efficiency, along with a no-insert control. The remaining volume was grown overnight at 30 °C. In the morning, plasmid DNA was extracted from the entire sample volume using a midiprep kit (Qiagen) according to the manufacturer’s instructions.

The αI^kinase^ library was prepared in two vector backbones. First, the BMX-H SH2 domain was used to replace the human SH2 domain in the BTK sequence. The BMX-H SH2 sequence was purchased from Twist Bioscience as a gene fragment, which was then amplified using oligos #468 and #469 (Table S2). The amplicon was ligated to the 5′ and 3′ arms of BTK using the ligation scheme above, and then the BamHI/XmaI fragment was ligated into the original digested pHIV vector. The αI^kinase^ library was purchased as an oligo pool from Twist Bioscience and amplified in four separate PCR reactions using the universal forward primer (oligo #432) and one reverse oligo (#433, #434, #435, or #436, Table S2). These reverse oligos included a three-nucleotide barcode after the stop codon and the BamHI restriction site. Two 5′ constant sequences were then prepared by amplifying either the human BTK sequence or the BMX-H SH2 chimera, both with oligos #437 and #275, but using their respective templates. These constant sequences were then ligated to the barcoded, amplified αI^kinase^ library by matching oligos #433 (barcode TTC) and #434 (barcode CCT) to the human SH2 domain and oligos #435 (barcode GGA) and #436 (barcode AAG) to the BMX-H SH2 domain. Ligation reactions were cloned into the pHIV backbone as above, sample was purified using a Midiprep kit (Qiagen), and then the resulting plasmid libraries were pooled prior to lentivirus preparation.

### Lentivirus preparation and titering

Lentiviruses were prepared as previously described^28^. Briefly, HEK293FT cells were seeded at a density of 250,000 cells/mL in 5 mL in 6-cm dishes in culture media (day 1). The next day (day 2), cells were transfected with a mixture of transfer plasmid containing the SH2-BTK chimeras or the αI^kinase^ chimeras (modified from pHIV-EGFP, addgene #21373, 5 µg) and two packaging plasmids (pMDG.2, #12259, 1.25 µg and psPAX2, #12260, 3.75 µg) in 500 µL of opti-mem (Thermo Fisher) using lipofectamine LTX (Thermo Fisher) according to the manufacturer’s instructions. The following morning (day 3) cells were re-fed. The following day (day 4) 5 mL of viral supernatant was harvested and stored at 4 °C, and the cells were re-fed. The following day (day 5) an additional 5 mL of viral supernatant was harvested and combined with the previous day’s aliquot for a total of 10 mL of viral supernatant. This was then concentrated using Lenti-X concentrator (Takara Bio) according to the manufacturer’s instructions, and the pellet from this concentration was resuspended in 1 mL of RPMI + 5% FBS, aliquoted, flash frozen in liquid nitrogen, and stored at –80 °C.

Lentiviruses were titered by thawing an aliquot of virus and adding 0, 5, 10, 20, 40, or 80 µL of virus to 500 µL ITK-deficient Jurkat T cells or BTK-deficient Ramos B cells at 500,000 cells / mL in a 24 well plate, along with 4 µg / mL polybrene. The following day, cells were centrifuged (300 g for 5 min) and resuspended in 1 mL of fresh media, then plated onto a 96-well plate. Cells were grown for two additional days and then the plate was analyzed for GFP fluorescence using an Attune flow cytometer (Thermo Fisher) equipped with a 96-well plate autosampler. The fraction of GFP positive cells as a function of concentration was fit to the standard binding isotherm to obtain titer values.

### Cell culture

All cells used in this study were cultured in a humidified incubator at 37 °C with 5% CO2. BTK-deficient Ramos B cells^28^ were cultured in RPMI (ThermoFisher) containing 10% fetal bovine serum (ThermoFisher) and 1x Glutamax (ThermoFisher). ITK-deficient Jurkat cells^29^ were cultured in RPMI containing 5% fetal bovine serum and 1x Glutamax. HEK293FT packaging cells, used for lentivirus preparation, were cultured in DMEM (ThermoFisher) containing 10% fetal bovine serum and 1x Glutamax.

### High-throughput assays

High-throughput experiments were performed as described in ^Ref. 28^. Briefly, ITK-deficient Jurkat T cells or BTK-deficient Ramos B cells were plated in 10 mL at a density of 500,000 cells / mL in 10 cm dishes in triplicates or quadruplicates. The cells were then transduced on day 1 with titered lentiviral libraries, adding a volume of virus such that at most 25% of the cells were GFP positive (corresponding to < 3% of cells having a multiplicity of infection > 1) and 10 µg / mL polybrene.

The following morning (day 2) cells were re-fed by centrifugation at 300 g for 5 min followed by resuspension in fresh media, with a 2x dilution, and re-plated. Cells were allowed to recover for one additional day (day 3). On day 4, Ramos cells were stimulated in the evening with 4 µg / mL antibody targeting IgM (I2386, Sigma-Aldrich).

On day 5, Ramos or Jurkat cells were stained with antibody targeting CD69 as follows.

Cells were concentrated by centrifugation at 300 g for 5 min, then resuspended in 200 µL of cell- staining buffer (phosphate-buffered saline containing 10% FBS and 0.05% w / v sodium azide) containing a 1:20 dilution of PerCP-Cy5.5 conjugated antibody targeting CD69 (Cell Signaling Technology #28633). Cells were then incubated on ice for 30 min in the dark. Following this incubation, cells were centrifuged again and washed with 500 µL cell-staining buffer, then centrifuged and resuspended in 1 mL of cell-staining buffer for sorting. After this staining protocol, 100 µL (10%) of sample was removed as input and kept on ice during sorting. Cells were sorted for a GFP-positive and CD69-positive population on a SH800 Cell Sorter (Sony) or FACSAria Fusion (BD Biosciences).

Gating was performed as follows. All cells were first gated based on forward and side scatter for a healthy population, generally 80-90% of the total cell number. To set the gates for the transduced (GFP-positive) and CD69 (PerCP-Cy5.5) positive populations, negative controls that were untransduced (GFP-negative) or unstained (PerCP-Cy5.5-negative) were used. Gates were set such that fewer than 0.1% negative-control cells fell within the positive gates. The double-positive population was then sorted for each library. In the case of Jurkat and Ramos cells, this double-positive population was approximately 5% of the total number of cells. Ramos cells required stimulatory antibody targeting IgM to express enough CD69 to sort 5% double- positive cells, while Jurkat cells did not require any stimulation. Each experiment sorted at least 200x the number of cells as variants in the library.

### Library preparation and sequencing

Following sorting, input and sorted samples were mixed with 1 mL of TRI reagent (Sigma-Aldrich), concentrating the sorted sample by centrifugation (300 g, 5 min) if the volume exceeded 200 µL. RNA was extracted according to the manufacturer’s protocol, except that after the aqueous phase was transferred to a new tube, it was mixed 1:1 with chloroform and centrifuged at 21,000 g for 10 min at 4 °C as an additional wash step. 4 µL of linear acrylamide (Thermo Fisher) was added as carrier.

RNA-seq libraries were prepared from TRI reagent-extracted samples as follows, beginning with the reverse-transcription. Following precipitation, RNA was resuspended in water containing 5 µM RT primer (oligo #296, Table S2). The mixture was heated to 65 °C for 5 min, then snap cooled on ice. The RNA–oligo mixture was then mixed with 1x First-Strand Buffer (Thermo Fisher), 0.5 mM deoxynucleotide triphosphates, 10 mM DTT, 1 µL SuperaseIn (Thermo Fisher), and 0.5 µL Superscript III (Thermo Fisher). The final reaction mixture was incubated at 50 °C for 1 h. Followed this incubation, RNA was hydrolyzed by addition of 5 µL of 1 M NaOH and incubation at 90 °C for 10 min. The mixture was then neutralized with 25 µL of 1 M HEPES, pH 7.4, and desalted using a Micro Bio-Spin P-30 tris column (BioRad), eluting the cDNA in 60 µL.

The cDNA was PCR amplified in two rounds. The first-round PCR added the priming regions for Illumina sequencing but did not add the barcodes (using oligos #278, #279 for the SH2 library and oligos #438 and #439 for the αI^kinase^ library, Table S2). The second-round PCR used primers that annealed to the Illumina priming regions, adding the flow-cell hybridizing sequences and barcodes (using primers BC_p5_1, BC_p5_2, BC_p5_3, BC_p5_4, BC_p5_5, BC_p5_6, BC_p5_7, BC_p5_8, BC_p7_1, BC_p7_2, BC_p7_3, BC_p7_4, BC_p7_5, BC_p7_6, BC_p7_7, BC_p7_8, BC_p7_9, BC_p7_10, BC_p7_11, and BC_p7_12, Table S2). Amplified libraries were gel-purified on a 2% agarose gel, extracted with a gel-extraction kit (Qiagen), pooled at equal molar based on the band intensities, and then concentrated using a Clean and Concentrator kit (Zymo). Libraries were sequenced using a MiSeq sequencer and V2 chemistry with paired-end 150 by 150 bp reads.

### Data analysis

Quantification of fitness from sequencing data was performed as previously described^28^. Briefly, Fastq files from MiSeq runs were aligned to the Fasta files containing the full sequences of each variant using Kallisto^30^ to generate read counts for each variant. A read cutoff of 50 reads was applied to the input libraries such that any variant not passing this threshold was discarded. Next, the unnormalized scores were calculated by dividing the number of reads in the sorted dataset by the number of reads in the input dataset and taking the base-10 logarithm. These unnormalized scores were normalized by subtracting the mean of the human BTK SH2 fitness scores, according to equation 1:

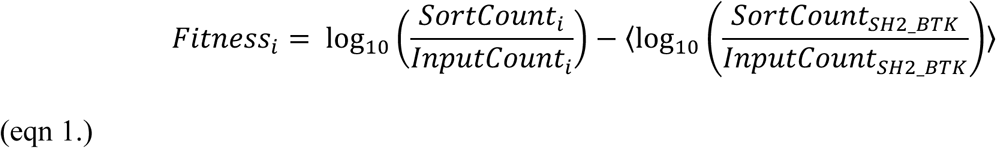

In equation 1, *Fitnessi* denotes the fitness score for a particular variant *i*. Because each sequencing library contained synonymous sequences of the human BTK sequence (rather than one human BTK sequence), fitness scores were calculated by subtracting the mean of these synonymous sequences.

Code to generate SH2-domain and αI^kinase^ libraries, perform ancestral-sequence reconstruction, and analyze RNA-seq libraries was written using R^Ref. 31^ and Python and is available on Github (https://github.com/timeisen/MutagenesisPlotCode and https://github.com/timeisen/SH2s).

## Supplemental Figure Legends

**Figure S1.**
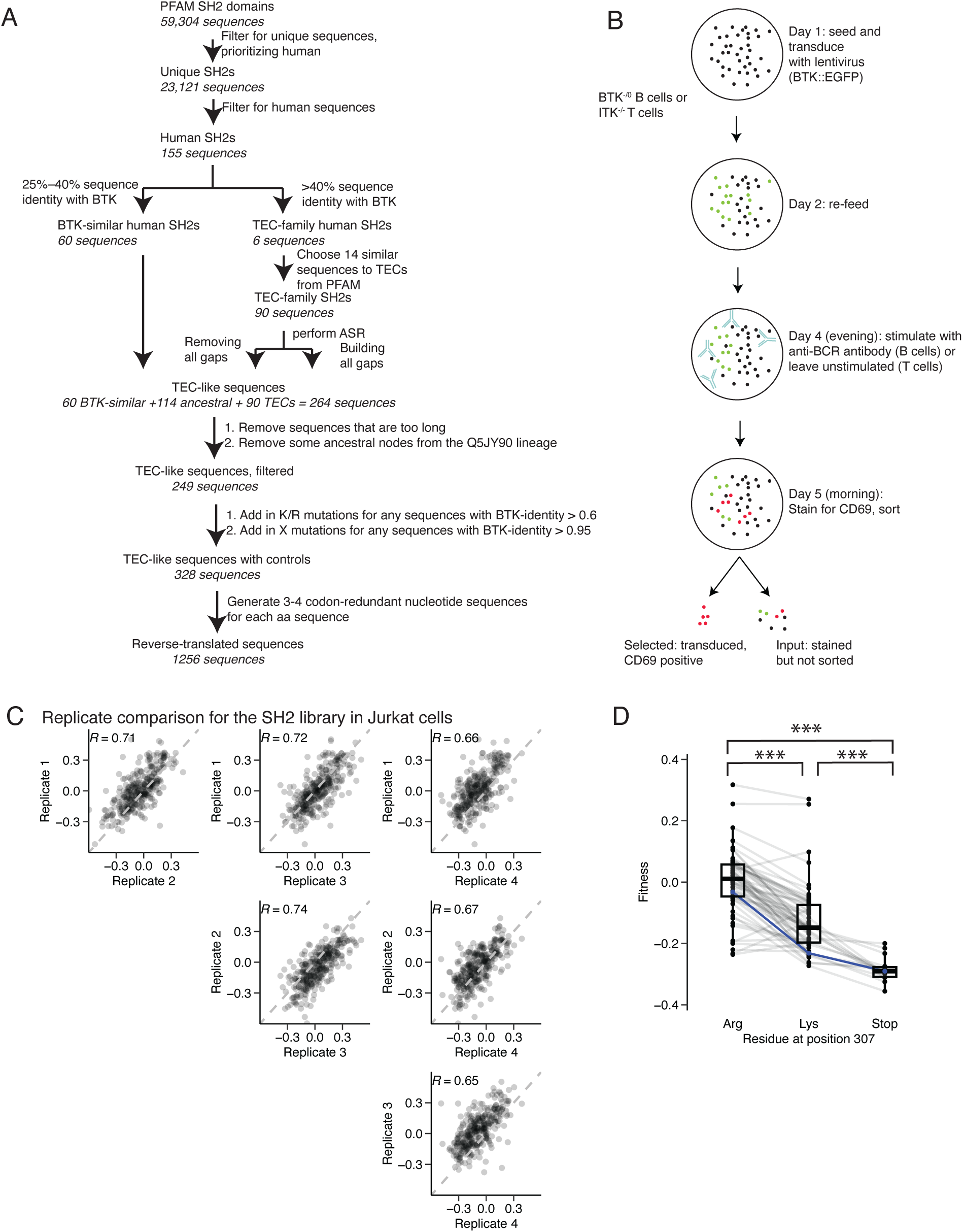
Design of the SH2-substitution experiment. a. Flow chart describing the computational steps to choose SH2 domains to substitute in human BTK. b. Schematic of the transduction and selection. ITK-deficient Jurkat cells or BTK-deficient Ramos cells were transduced with a library of viruses containing BTK or chimeras, an IRES, and EGFP. After 3 days of culture, Ramos cells were stimulated with anti-IgM stimulatory antibody at 4 µg / mL. The next day, cells were stained for CD69 and sorted for both EGFP (marking the transduced population) and CD69. An input fraction, taken just before sorting, was used to determine enrichments. c. Pairwise comparisons between four biological replicates for the n = 328 SH2 domains. The indicated R is the Pearson correlation between the two samples, and the dashed line indicates the y = x diagonal. d. Fitness scores from mutations to Arg 307, the phosphotyrosine-interacting residue. SH2 domains derived from the Tec kinases or ancestral-sequence reconstruction were mutated to replace the Arg residue at the equivalent position to 307 in human BTK with either lysine (62 sequences) or a stop codon (17 sequences). The fitness scores from each domain in all three contexts are shown, with lines connecting the same domain (because not all sequences were mutated with a stop codon, lines do not always span all three groups). The blue line is the human BTK SH2 domain. A box-and-whiskers plot is overlayed with the line (median), box (first and third quartiles) and whiskers (1.5x the interquartile range). Asterisks denote ***P < 0.001 with a one-way ANOVA with Tukey’s multiple-comparison tests.

**Figure S2.**
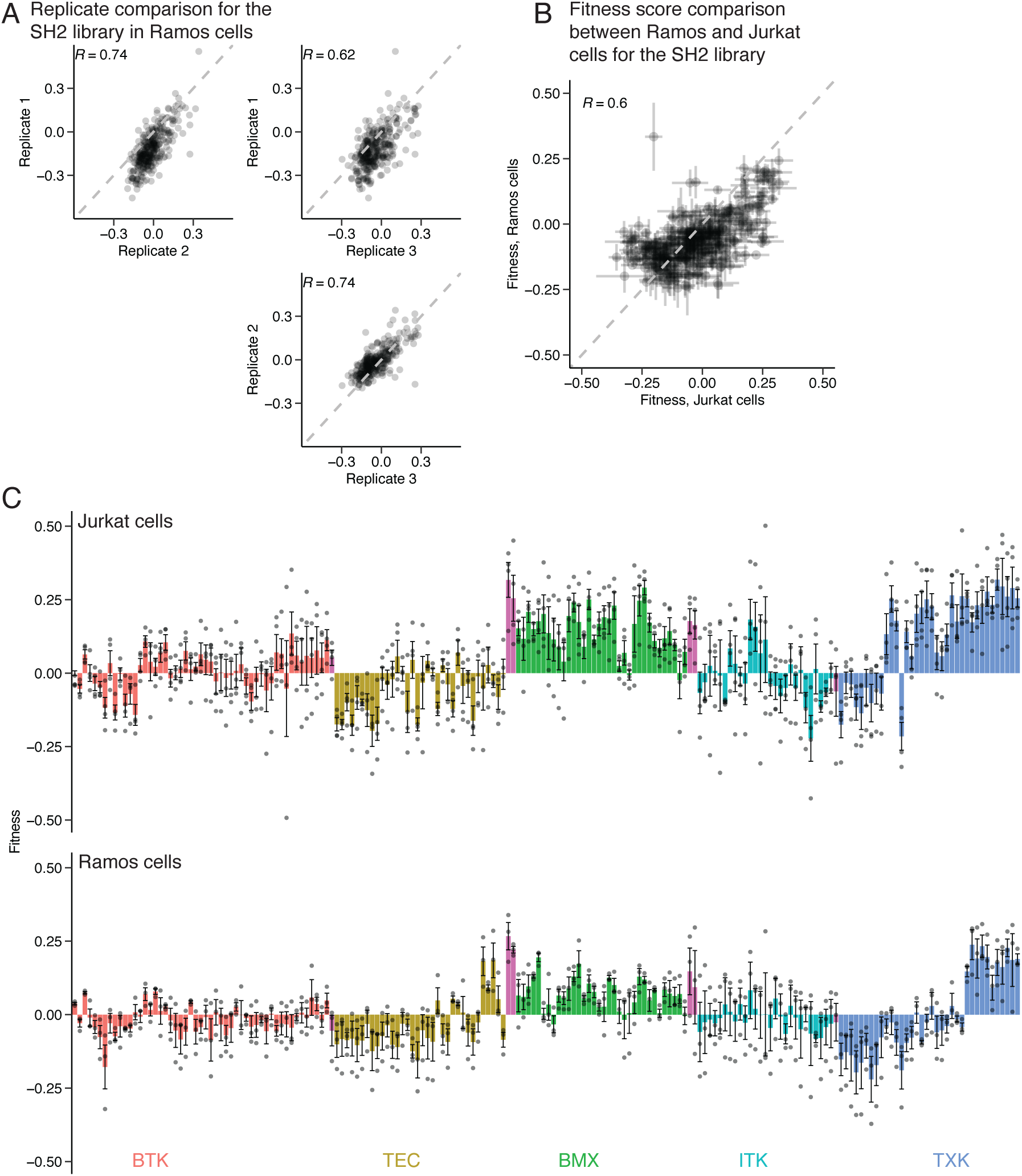
Comparison between fitness scores in Jurkat T cells and Ramos B cells. a. Pairwise comparisons between three biological replicates for the n = 328 SH2 domain chimeras in the Ramos assay. The indicated R is the Pearson correlation between the two samples, and the dashed line indicates the y = x diagonal. b. Comparison between the mean fitness values for each of the n = 328 SH2 domain chimeras in the Jurkat and Ramos assays. The fitness values are the mean of the biological replicates, and the error bars represent the standard error of the mean. c. Comparison of individual SH2 sequences between Jurkat and Ramos cells, as in Figure 1C.

**Figure S3.**
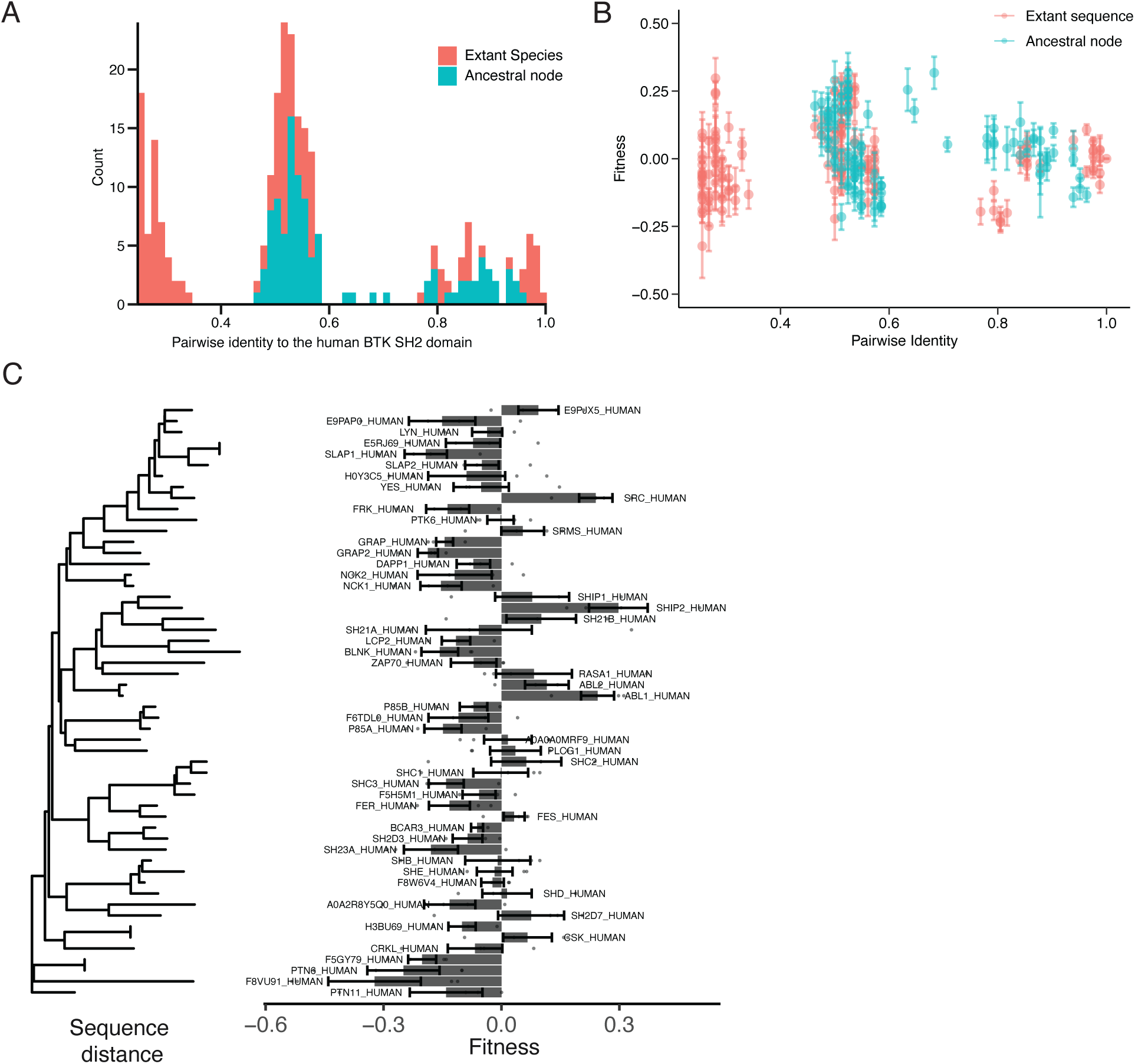
Ancestral and extant sequence comparisons. a. A histogram of the pairwise identities of the SH2 domains included in the library, colored by whether the domains are derived from pre-existing proteins or generated using ancestral-sequence reconstruction. Pairwise identities are with respect to the human BTK SH2 domain sequence. Note that the only domains with pairwise identities in between those of the Tec-kinase group (∼50%) and the BTK group (90%) are ancestral, because ancestral-sequence reconstruction allows the generation of sequences with intermediate pairwise identity. The most distant sequences, which consist of SH2 domains from human proteins other than the Tec kinases, were not used for ancestral-sequence reconstruction. b. Relationship between the fitness score of each SH2 domain and its pairwise identity to the human BTK SH2 domain. c. Fitness scores for 54 human SH2-domain sequences with between 25% and 40% identity to the human BTK SH2 domain. Sequences are arranged on a phylogenetic tree, with error bars representing the standard error.

**Figure S4.**
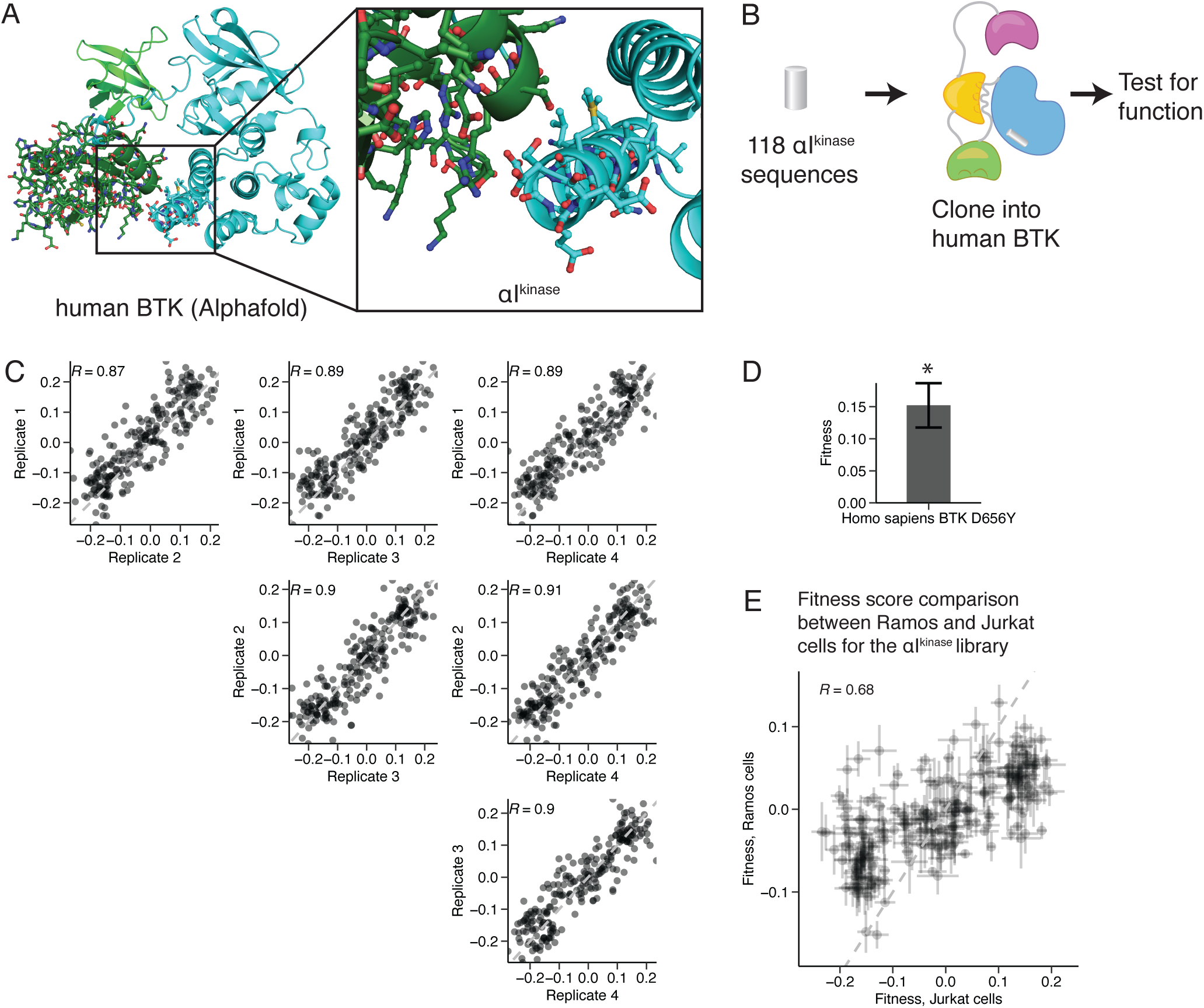
Substitution of αI^kinase^. a. Alphafold^20^ predicted structure of human BTK, with inset showing αI^kinase^. b. Schematic of the αI^kinase^ swapping experiment. c. Replicate correlations for the fitness scores for the 236 proteins assayed (118 αI^kinase^ variants in two genetic backgrounds). The indicated R is the Pearson correlation between the two samples, and the dashed line indicates the y = x diagonal. d. The D656Y substitution increase fitness in human BTK. An αI^kinase^ sequence containing this single substitution was included in the αI^kinase^ library and assayed in Jurkat cells. Error bars represent standard error of the mean. The asterisk indicates that this substitution is significantly different than 0 with P < 0.05 for the n = 4 replicates, t-test. e. Comparison between the mean fitness values in Jurkat and Ramos cells for each of the 118 αI^kinase^ chimeras in both the human BTK SH2 and BMX-H SH2 genetic backgrounds (n = 236 total). Error bars represent standard error of the mean.

**Figure S5.**
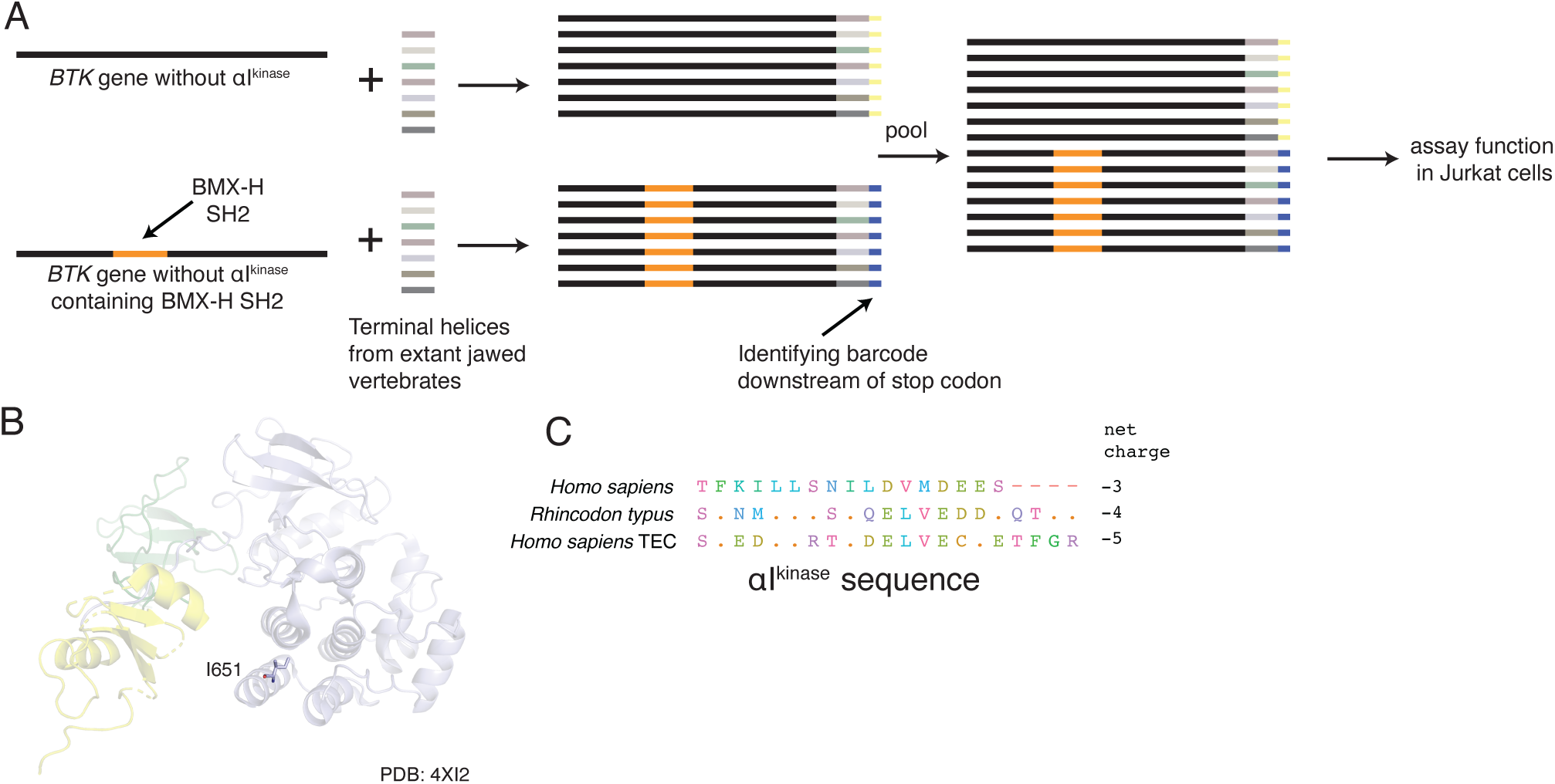
Construction of the αI^kinase^ library in two genetic backgrounds. a. Schematic of the library cloning for the αI^kinase^ swapping experiment. A library of αI^kinase^ sequences was used to replace the αI^kinase^ sequence in either human BTK or the BMX-H chimera. These two sets of libraries were then barcoded with a 3-nucleotide barcode after the stop codon, and pooled. A single sequencing read was used to determine the identity of αI^kinase^ and the barcode, which indicated whether the SH2 domain was BMX-H or human BTK. b. Crystal structure of mouse Btk, highlighting the position of Ile 651. PDB: 4XI2^Ref. 8^. c. αI^kinase^ sequences of human BTK, *R. typus* BTK, and human TEC kinase. The net charge across the sequence (the number of glutamates or aspartates subtracted from the number of lysines or arginines) is shown at the right.

**Table S1. Fitness scores and sequences of all SH2 domains examined in this study in both cell lines.**

**Table S2. Oligonucleotides used in this study.**

